# Influence of pseudo-green water on growth, colouration, and survival in European lobster (*Homarus gammarus*) larvae

**DOI:** 10.1101/2023.08.30.555547

**Authors:** Matt E Bell, Julie O Grady, Paula C Domech, Erik R Frontaura, James Hinchcliffe, Simon J Davies, Stephen McCusker, Alex H L Wan

**Affiliations:** Aquaculture and Nutrition Research Unit (ANRU), Carna Research Station, School of Natural Sciences and Ryan Institute, University Galway, Carna, Co. Galway, Ireland, H91 V8Y1; Aquaculture and Nutrition Research Unit (ANRU), School of Natural Sciences and Ryan Institute, Annex building, University Galway, Carna, Co. Galway, Ireland, H91 TK33; Department of Biological and Environmental Sciences, The Swedish Mariculture Research Centre (SWEMARC), University of Gothenburg, Gothenburg, Sweden

**Keywords:** Green water, Larval Culture, Hatchery, Carotenoids, Crustaceans, Decapods

## Abstract

With the depletion of European lobster (*Homarus gammarus*) wild stocks from historic overexploitation, the attention has turned towards restocking, stock enhancement programs and farming potential. The early life larval stages of the European lobster present a major barrier to hatchery production efforts due to low survival rates. Green water techniques are rapidly growing due to their ability to improve nutrition quality. This study explores the potential benefits of using pseudo-green water. This is determined by adding microalgae to the rearing environment of larvae by analysing the effects on growth and survival. The commonly used algal species: *Tetraselmis suecica* (TS), *Dunaliella salina* (DS), and *Nannochloropsis* sp. (N) were added as monoculture or mixture treatments to a controlled European lobster larvae-rearing environment over thirteen days. The trial consisted of seven permutations of algal treatments (TS, DS, N, TS+DS, TS+N, DS+N, and TS+DS+N) and one control group, with clear water conditions using length measurements, survival rates and colouration to determine treatment impacts at the end of the trial. Results indicate that mixtures of algal species outperformed monocultures and clear water regarding growth performance. *Nannochloropsis* sp. is of significant benefit to growth performance. Algal mixes containing *Nannochloropsis* sp. resulted in significantly better growth than *D*. *salina* monoculture treatment. There were no significant differences in survival rates between treatment groups.

## 1 Introduction

Lobsters act as keystone species in benthic communities, employing top-down predatory control on complex food webs. The ubiquitous reduction in lobster species biomass in terms of catch has had structural and functional impacts on their ecosystems (Eddy et al., 2014). It is well known that current wild European lobster (*Homarus gammarus*, Linnaeus, 1758) stocks are under immense pressure due to several factors, primarily overfishing and climate change, including effects on their physiology, metabolism, and bioenergetics and rising sea temperatures. Furthermore, specific abiotic concerns include the potential electromagnetic effect of offshore wind energy installations on natural larval recruitment and survival (Harsanyi et al., 2022). The European lobster is a commercially valuable marine crustacean for seafood whilst supporting fishing industries throughout European coastal waters. For example, when combining the UK, the Republic of Ireland, Sweden and Norway, the industry is worth €40 million in 2021 (Hinchcliffe et al., 2022). The FAO in 2021 reported that over 4,800 tonnes (live weight) of European lobster were landed (FAO, 2023). To safeguard current and future populations, considerable effort has been implemented into developing aquaculture technologies and hatchery production intervention for restocking and stock enhancement purposes in several European states (Hinchcliffe et al., 2022).

The early life history of European lobsters is characterised by four distinct larval stages. During the zoea I-III stages, they are planktonic and omnivorous (Rötzer & Haug, 2015). At the megalopa stage IV it is marked by metamorphosis (Charmantier et al., 1991) and the transition to a more benthic lifestyle before undergoing complete settlement during the early benthic settling phase (EBP) as juveniles (Linnane et al., 2000). A significant issue hindering hatchery production is the low survival rate through the planktonic larval phases, which rarely exceeds 20 % and is considered a recruitment bottleneck within even wild lobsters caused by predation (Addison and Bannister, 1994; Daniels et al., 2010; Ellis et al., 2015; Goncalves et al., 2023; Powell et al., 2017). Poor nutrition within artificial rearing is linked to stress and high mortality (Schoo et al., 2014), resulting in cannibalistic tendencies (Powell et al., 2017; Romano and Zeng, 2017).

The pseudo-green water techniques combine ‘clear water’ and ‘green water’ techniques by adding microalgae only during the most critical rearing period – larval rearing (Papandroulakis et al., 2001). Pseudo-green water techniques have demonstrated the benefits of algal addition to rearing systems (Palmer et al., 2007). These include improved water quality and the antimicrobial potential to mitigate pathogens (Brown, 2002; Palmer et al., 2007). This was identified in American lobster (*Homarus americanus*), where the authors hypothesised that green algae enrichment directly increases the nutritional value as part of the surrounding aqueous medium (Haché et al., 2017). It is possible that the green water is acting as a probiotic, whereby it is outcompeting harmful pathogens, improves background contrast to highlight motile prey items, and/or stimulate predator behaviour interactions.

Several different algal species are routinely employed as feed enrichment or green water in fish, echinoderm, and decapod hatcheries. For example, as observed in the rearing of turbot (*Scophthalmus maximus*), when *Tetraselmis suecica* (Prasinophyte) and *Isochrysis galbana* microalgae were added it increases marine fish larval survival rates to 63.5% (Pyanov, 2021). Another commercially important algae is *Dunaliella salina* (chlorophyte), due to its biosynthesis of β-carotene, an important functional carotenoid and a precursor to vitamin A (Wade et al., 2017). While *Nannochloropsis sp*. (Eustigmatophyte) is valued for its protein content (i.e., complete essential amino acid profile) and fatty acid profile. In particular, the high long-chain fatty acids of the omega-3 family, e.g., EPA (20:5n3) at 3 % dry weight and 17% of the total overall fatty acid profile for the medium 50-50 and 13.7 % of the total overall fatty acid profile for the 100*-0 medium (González-López et al., 2013).

These aspects warrant more information for applications in lobster larval rearing and the concept of using either single algal strains or mixtures to assess synergistic attributes in the culture system. Consequently, an experiment was conducted using the pseudo-green water technique to evaluate the feasibility of varying microalgal additions to the rearing environment of European lobster larvae on growth performance, body colouration, and survival to zoea stage III of development. The effects of mono, dual, and mixed cultures of *T. suecica*, *D. salina,* and *Nannochloropsis* sp. commonly used in green water culture was assessed for their performance to enhance survival and growth at the zoea stages of lobsters.

## 2. Materials and Methods

### 2.1. Lobsters and experimental design

Larvae were obtained from a berried European lobster (*Homarus gammarus*) female. Stage I larvae were collected within two days of release. Ten larvae individuals were randomly allocated into each thirty-two 1 L polycarbonate bottles. The trial study consisted of eight culture treatments; seven treatments were different permutations of pseudo-green water: [1] *Tetraselmis suecica* (T); [2] *Dunaliella salina* (D); [3] *Nannochloropsis* sp. (N); [4] *T. suecica* & *D. salina* (1:1, T + D); [5] *T. suecica* & *N.* sp. (1:1, T + N); [6] *D. salina* & *N*. sp. (1:1, D + N); [7] *T. suecica, D. salina* & *N.* sp. (1:1:1, T + D + N). The eighth treatment was the control which was clear water with no algal addition (C). Each treatment was tested in four replicate culture vessels (n=4). Bottles were filled with pasteurised seawater and provided with constant gentle aeration to add oxygen and create turbulence to ensure the lobster larvae remained suspended. Lighting to the bottles were set at a 12:12 photoperiod (08:00 am to 20:00 pm). Temperature was 19.09 ± 1.09 °C (SD) during the trial period. Lobsters were fed daily with artemia enriched (Easy dry SELCO, INVE, Dendermonde, Belgium) at ∼60 artemia L^-1^. A complete water change was undertaken every four days.

### 2.2. Algal culture

Microalgae strains of *Tetraselmis suecica, Dunaliella salina*, and *Nannochloropsis* sp. was selected for the trial with different nutrient composition (Table 1). Algal cultures were grown from stocks at Carna Research Station, Ryan Institute, University of Galway, Carna, Ireland. Cultures were enriched with F/2 nutrient media (CELL-HI F2P, Varicon Aqua Solutions, Worcestershire, UK) and were grown in constant fluorescent light (Osram de lux, Munich, Germany). Algae cultures were added at day zero and post-water changes (i.e., days 4, 8 and 12). The desired volumes of algae culture were calculated based on cell counts carried out on the day of inoculation. A subsample of each algae culture was fixed and stained in lugol solution (1:5), and cell counting was undertaken using a haemocytometer. The amount of algae culture used was calculated based on the desired cell count of ^∼^1,000,000 L^-1^ in the culture vessel.

**Table 1:** Proximate composition and fatty acid composition of algal species used in the present study: *Tetraselmis suecica*, *Dunaliella salina,* and *Nannochloropsis* sp.

|  | *Microalgae species* |  |  |
| --- | --- | --- | --- |
|  | *Tetraselmis suecica* | *Dunaliella salina* | *Nannochloropsis* sp. |
| *Proximate composition<sup>a</sup>* |  |  |  |
| Protein | 41 | 57 | 55 |
| Lipid | 24 | 6 | 37-60 |
| Carbohydrate | 12 | 32 | 18 |
| *Fatty acid profile<sup>b</sup>* |  |  |  |
| ΣSFA | 33.02 - 36.68 | 26.24 - 39.10 | 21.23 - 50.87 |
| ΣMUFA | 17.27 - 35.92 | 33.39 - 46.05 | 31.60 - 52.56 |
| ΣPUFA | 31.06 - 46.05 | 27.51 - 34.84 | 6.66 - 41.70 |
| *References* | Fernández-Reiriz et al., 1989; Guzmán et al., 2010 | Gastelum-Franco et al., 2021; Spolaore et al., 2006 | Brennan & Regan, 2020; Ma et al., 2014; Peltomaa et al., 2017; Recht et al., 2012; |

<sup>a</sup>% dry weight; <sup>b</sup>%TFA, % of total fatty acid content. SAT, saturated fatty acids; MUFA, monounsaturated fatty acids; PUFA, polyunsaturated fatty acids.

### 2.3. Morphometric measurements

On day 0 and 13 of the study, images of the lobster larvae were taken using a digital single-lens reflex camera (D810, Nikon, Tokyo, Japan) using a 110mm lens (1:1, Sigma Corporation, Setagaya, Japan) with a ring flash system (Commander Kit R1C1, Nikon, Tokyo, Japan). The images were analysed for morphological parameters, survival, and colouration. A scale bar and a colour panel (Konica Minolta, Chiyoda City, Tokyo, Japan) was included in each image that was taken for calibration. The following indices were calculated:

**Weight gain, %** = (Final lobster weight (g) – Initial lobster weight (g)) *100

**Carapace length increase**; **CL**, % = (Final lobster carapace length (mm) - Initial lobster carapace length (mm)) *100

**Total length increase**; **TL**, % = (Final lobster total length (mm)– Initial lobster total length (mm)) *100

**Survival**; % = (Final lobster individual count – Initial lobster individual count) *100

### 2.4. Image analysis

Image analysis of *H. gammarus* larvae was conducted using Fiji v2.13.1 image analysis software (Schindelin et al., 2012). The in-image scale bar calibrated the total length (mm) and carapace length (mm). Prior to the measurement of the lobster colouration, images were processed, and colour was calibrated using an in-image reference colour tile through Photoshop CS5 (Tlusty, 2005). Colour analysis was conducted in Fiji (Schindelin et al., 2012) to obtain R G B values for points on the claw, cephalothorax, and abdomen. RGB values were converted to *L* a* b\** values (CIE 2004). *L*a*b\** values best represent colours as seen by the human eye. *L*\* indicates perceptual lightness, 0 darkness/100 lightness; *a*\*+, redness/ *a*\*-, greenness; and *b*\*+ blueness/ *b*\*-yellowness.

ΔE* is an empirical metric expressing the variation between two colour values based on a scale from 0 – 100; values less than one are unperceivable by the human eye. Values in the range of 11 – 49 describe colours that are more similar than opposite (Daniel, 2021). The total colour difference, ΔE*, between each green water treatment and the control group (no algae) was calculated using the following formula:

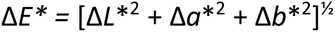

### 2.5. Statistical analysis

The effect of microalgae composition on lobster larval growth parameters were initially tested by one-way analysis of variance (ANOVA) on morphometric and colour parameters. Datasets were tested for normal distribution and homogeneity of variance using Shapiro-Wilk and Levene tests, respectively. Non-normal distribution datasets were subsequently analysed using the non-parametric Kruskal-Wallis test. To discern statistical differences between algal treatments, the post hoc Tukey test was used. The contributing influence of each algal species towards the larval performance was further assessed through a mixture design analysis of variance model using Design Expert 7.0.0 (Stat-Ease, 2005).

## 3. Results and Discussion

Captive rearing culture of European lobster (*H. gammarus*) larvae has presented significant challenges over the years, and ultimately survival ratios from pelagic stage I larvae to benthic stage IV+ have been low. Past studies have reported variable survival rates from 3-15 %, with the present study attaining a comparable survival rate of 12.5-20 % (Ellis et al., 2015; Önal & Baki, 2021). Many researchers have encountered similar bottlenecks and it is generally accepted that the somewhat indiscriminate use of microalgae species in larval lobster culturing may lead to inevitable nutrient deficiencies, such as key polyunsaturated fatty acids (PUFAs), vitamins such as ascorbic acid and important pigments such as lutein, astaxanthin and canthaxanthin (Becker, 2013; Guedes & Malcata, 2012).

The present study employed a pseudo-green water technique, using the microalgal species *Tetraselmis suecica, Dunaliella salina,* and *Nannochloropsis* sp., as both monocultures and mixtures. The goal was to determine whether this approach significantly impacted the growth and survival of European lobster pelagic larvae. The study was conducted over 13 days and all larvae had reached stage III, where cannibalism typically starts to occur.

Several previous investigations demonstrate the proven benefit of providing algal mixtures over monospecific cultures, e.g., in juvenile clams (*Tapes semidecussata*) and Pacific white shrimp (*Litopenaeus vannamei*) (Becker, 2013; Piña et al., 2006). Furthermore, a dietary source of mixed microalgal has been shown to outperform monoalgal cultures in the pullet carpet shell clam (*Venerupis pullastra*), hard clam (*Mercenaria mercenaria*), American oyster (*Crassostrea virginica*) hatcheries (Albentosa et al., 1993; Epifanio, 1979), and in sea urchin (Paracentrotus *lividus*) larviculture (Gomes et al., 2021).

### 3.1. Growth performance

There was no statistical difference (p > 0.05) between the length measurements of the treatment groups on day 0 (Table 2). Therefore, this did not affect the outcome of day 13 length measurements: hence, analyses on the mean length measurements of day 13. Both total and carapace lengths from all mixture groups (T + D; T + N; D + N; T + D + N) were greater than that of the control group (C), though not significantly. However, there were significant differences (p < 0.05) for carapace lengths and percentage gains. Excluding the control, D resulted in the smallest total (12.97 ± 0.59 mm) and carapace length (5.13 ± 0.26 mm), compared to T + N, which had the highest total (14.36 ± 0.43 mm) and carapace length (6.08 ± 0.93 mm), equating to a 10.7 % and an 18.5 % increase in growth respectively. Past research studies reveal that dietary lipid content provides an important contribution to European lobster larvae nutritional profiles for energy and metabolic purposes (Becker, 2013; Goncalves et al., 2022). *Nannochloropsis* sp. has a considerably higher total lipid content (37-60%) when compared to *D. salina* (6%), which likely influenced the significant differences in growth results between treatment groups (Table 1).

**Table 2:**
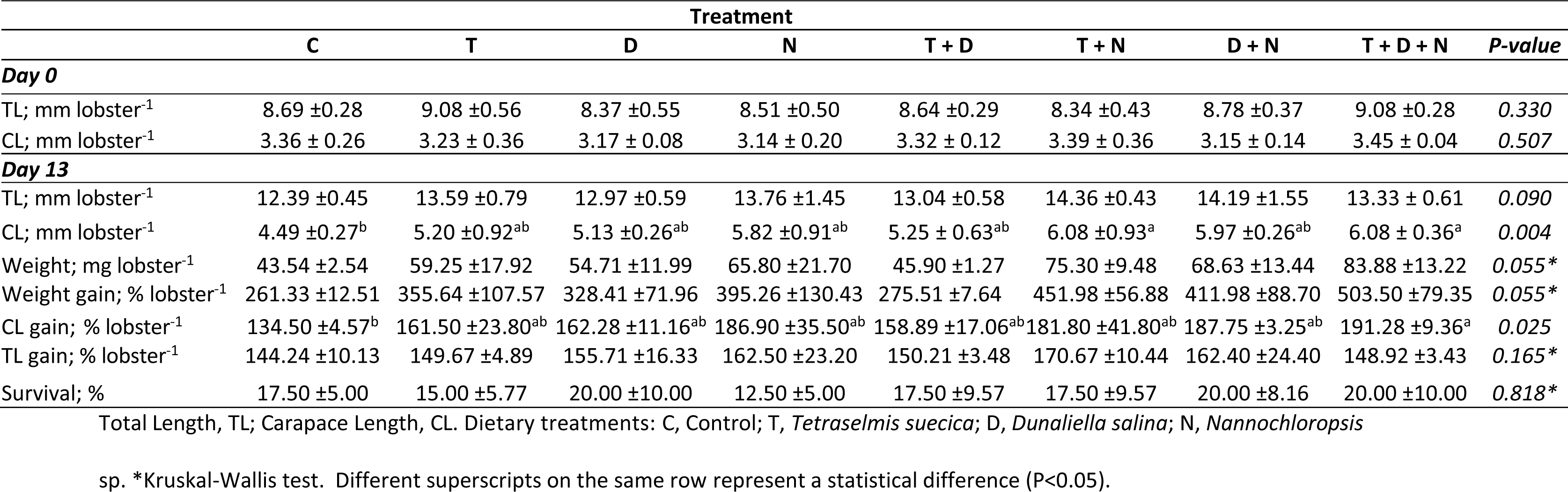
The effect of pseudo-green water on European lobster (*Homarus gammarus*) rearing has on morphological indices, growth performance, and survival over 13 days post hatching (n=4, ± SD).

|  | Treatment |  |  |  |  |  |  |  |  |
| --- | --- | --- | --- | --- | --- | --- | --- | --- | --- |
|  | C | T | D | N | T + D | T + N | D + N | T + D + N | *P-value* |
| **Day 0** |  |  |  |  |  |  |  |  |  |
| TL; mm lobster<sup>-1</sup> | 8.69 ±0.28 | 9.08 ±0.56 | 8.37 ±0.55 | 8.51 ±0.50 | 8.64 ±0.29 | 8.34 ±0.43 | 8.78 ±0.37 | 9.08 ±0.28 | 0.330 |
| CL; mm lobster<sup>-1</sup> | 3.36 ± 0.26 | 3.23 ± 0.36 | 3.17 ± 0.08 | 3.14 ± 0.20 | 3.32 ± 0.12 | 3.39 ± 0.36 | 3.15 ± 0.14 | 3.45 ± 0.04 | 0.507 |
| **Day 13** |  |  |  |  |  |  |  |  |  |
| TL; mm lobster<sup>-1</sup> | 12.39 ±0.45 | 13.59 ±0.79 | 12.97 ±0.59 | 13.76 ±1.45 | 13.04 ±0.58 | 14.36 ±0.43 | 14.19 ±1.55 | 13.33 ± 0.61 | 0.090 |
| CL; mm lobster<sup>-1</sup> | 4.49 ±0.27<sup>b</sup> | 5.20 ±0.92<sup>ab</sup> | 5.13 ±0.26<sup>ab</sup> | 5.82 ±0.91<sup>ab</sup> | 5.25 ± 0.63<sup>ab</sup> | 6.08 ±0.93<sup>a</sup> | 5.97 ±0.26<sup>ab</sup> | 6.08 ± 0.36<sup>a</sup> | 0.004 |
| Weight; mg lobster<sup>-1</sup> | 43.54 ±2.54 | 59.25 ±17.92 | 54.71 ±11.99 | 65.80 ±21.70 | 45.90 ±1.27 | 75.30 ±9.48 | 68.63 ±13.44 | 83.88 ±13.22 | 0.055* |
| Weight gain; % lobster<sup>-1</sup> | 261.33 ±12.51 | 355.64 ±107.57 | 328.41 ±71.96 | 395.26 ±130.43 | 275.51 ±7.64 | 451.98 ±56.88 | 411.98 ±88.70 | 503.50 ±79.35 | 0.055* |
| CL gain; % lobster<sup>-1</sup> | 134.50 ±4.57<sup>b</sup> | 161.50 ±23.80<sup>ab</sup> | 162.28 ±11.16<sup>ab</sup> | 186.90 ±35.50<sup>ab</sup> | 158.89 ±17.06<sup>ab</sup> | 181.80 ±41.80<sup>ab</sup> | 187.75 ±3.25<sup>ab</sup> | 191.28 ±9.36<sup>a</sup> | 0.025 |
| TL gain; % lobster<sup>-1</sup> | 144.24 ±10.13 | 149.67 ±4.89 | 155.71 ±16.33 | 162.50 ±23.20 | 150.21 ±3.48 | 170.67 ±10.44 | 162.40 ±24.40 | 148.92 ±3.43 | 0.165* |
| Survival; % | 17.50 ±5.00 | 15.00 ±5.77 | 20.00 ±10.00 | 12.50 ±5.00 | 17.50 ±9.57 | 17.50 ±9.57 | 20.00 ±8.16 | 20.00 ±10.00 | 0.818* |

Total Length, TL; Carapace Length, CL. Dietary treatments: C, Control; T, *Tetraselmis suecica*; D, *Dunaliella salina*; N, *Nannochloropsis*
sp. \*Kruskal-Wallis test. Different superscripts on the same row represent a statistical difference (P<0.05).

Dietary lipids are an important source of essential fatty acids (EFAs) which are essential for normal growth performance and survival in European lobster larvae (Rainuzzo et al., 1997; Goncalves et al., 2022; Becker, 2013). The benefit of lipids for early larval growth and development can also be seen in other crustacean species, such as the temperate spider crab (*Maja brachydactyla*, Andrés et al., 2010), spiny lobster (*Panulirus argus*, Perera et al., 2008), and American lobster (*Homarus americanus*, Thériault and Pernet, 2007).

*Nannochloropsis salina,* which contains a high level of the PUFA eicosapentaenoic acid (EPA; 20:5n-3), has been shown to improve growth performance and survival in the phyllosoma larvae of the tropical spiny lobster (*Panulirus homarus*) (Radhakrishnan et al., 2009). Results were attributed to improved water quality and enhanced *Artemia* enrichment through constant feeding on *N. salina*. In our study, water quality impacts were unlikely due to frequent water changes and short trial duration. Improved growth in mixture groups containing *Nannochloropsis* sp. could be attributed to the high PUFA content of *Nannochloropsis* sp. and supplementary nutritional benefits provided by the other algal species in the mixture.

Results from the mixture design on the influence of the algae on growth measurements demonstrate the inadequacy of morphometric parameters resulting from the presentation of monotype diets. As such, the lowest recorded final body lengths occurred under single algae treatments. This supports previous research on the benefits of mixed diets (Brown, 2002; Brown et al., 1997; Guedes & Malcata, 2012; Piña et al., 2006). This difference in effect could be the result of a single algal species not providing sufficient nutrients to meet the full requirements of the animal.

While mixed algal cultures produce a more complete nutrient composition, thus more likely to provide all the dietary needs of the growing larvae (Guedes & Malcata, 2012).

*Nannochloropsis* sp. was influential within the algal mixtures, and this is likely due to its high total lipids of 37-60 % dry weight and PUFAs of 41.7 % total fatty acid content, which is critical nutritional components for European lobster larvae (Brennan & Regan, 2020; Goncalves et al., 2022; Ma et al., 2014). Despite being the only significantly influential algal species, it was not effective as a monoculture, indicating that *T. suecica* and *D. salina* could be of some benefit when incorporated with *Nannochloropsis* sp.

### 3.2. Larval colouration

Carotenoids such as astaxanthin and β-carotene are directly responsible for numerous health benefits and critical metabolic adaptions in larval lobster development. For instance, these natural pigmenting agents are an important antioxidant source beyond their characteristic colouration properties (Maoka, 2011; Wade et al., 2017; Yeap et al., 2022). Examples of commonly utilised microalgae species that are rich in carotenoids include *T. suecica, I. galbana* and *D. salina* (Di Lena et al., 2018; Raposo et al., 2015). Under suitable conditions, European lobsters will develop the typical darker and bluer colouration, however, it has been shown that lobster colouration is also attributed to environmental habitat setting (Tlusty et al., 2009).

Results from statistical analysis on *L* a* b\** values yielded no significant differences (P>0.05) between dietary treatments for the claw and abdomen colour levels (Table 2). The only significant result (P<0.05) was *b\** for the cephalothorax (Table 2). Colour development plays a crucial role in lobster larvae survival rates. Studies carried out on the Caribbean spiny lobster (*Panilurus argus*) suggest a preferential shift to disruptive colour patterns in smaller individuals (<23 mm carapace length), compared to bicolour pigmentation in adult lobsters (Anderson et al., 2012). Only T + D (*ΔE\** = 14-15) and D + N (*ΔE\** = 14-16) presented the most uniform darker colouration for all parts measured, i.e., claw, cephalothorax, and abdomen (Table 3). Analogous to dietary feeds, the environment plays a significant role in developing pigmentation through factors such as background colouration (McLean, 2021). This would suggest that a mixture, particularly T + D and D + N, provides the best outcomes compared to mono treatments in delivering better survival and performance, i.e., higher pigmentation allows better crypsis. Further evidence to support the statement that mixed treatments provide beneficial optimisation can be seen with the *ΔE\** values. Mixed treatments produce more optimal colouration than mono treatments, for example, claw, D + N, T + N and T + D, compared to only treatment N, which insolently provided the highest colouration at 18.58 *ΔE\** (Table 3). The same is true for the cephalothorax: D + N and T + D, compared to T, once more providing the highest colouration value at 20.99 *ΔE\** (Table 3). This is also the case for the abdomen, with T + D + N, D + N and T + D providing beneficial colouration compared to only T and N from the monocultures (Table 3). It is evident from this that treatments containing D prove the most positive commercial outcome, and the mix D + N is the most effective for commercial value in colouration. Without using carotenoid extraction from end-trial larvae and algal cultures, it is difficult to determine the effect of algal carotenoid content on resulting larvae colouration. Further research is required to assess pigmentation development in response to green water treatment by direct qualitative and quantitative analysis.

**Table 3:** The influence of pseudo-green water on European lobster (*Homarus gammarus*) larvae shell colouration after 13 days post-hatching (claw, cephalothorax, and abdomen, n=4, ±SD).

| Treatment |  |  |  |  |  |  |  |  |  |
| --- | --- | --- | --- | --- | --- | --- | --- | --- | --- |
|  | C | T | D | N | T + D | T + N | D + N | T + D + N | P-values |
| Claw |  |  |  |  |  |  |  |  |  |
| *L*<sup>*</sup> | 42.89 ±4.28 | 45.70 ±12.13 | 50.08 ±9.25 | 55.50 ±4.12 | 41.20 ±0.36 | 52.31 ±0.86 | 54.71 ±14.45 | 47.48 ±13.81 | 0.481 |
| *a*<sup>*</sup> | 35.31 ±5.95 | 33.01 ±7.59 | 30.77 ±6.64 | 27.82 ±14.38 | 23.88 ±1.89 | 34.23 ±3.15 | 27.11 ±9.69 | 34.75 ±8.25 | 0.555<sup>*</sup> |
| *b*<sup>*</sup> | 23.78 ±5.51 | 27.58 ±12.61 | 26.85 ±5.64 | 35.19 ±3.68 | 14.92 ±0.20 | 35.40 ±0.53 | 32.35 ±12.45 | 31.12 ±16.12 | 0.292 |
| Δ*E*<sup>*</sup> | - | 5.25 | 9.03 | 18.58 | 14.57 | 14.98 | 16.75 | 8.67 |  |
| Cephalothorax |  |  |  |  |  |  |  |  |  |
| *L*<sup>*</sup> | 31.91 ±7.39 | 20.44 ±3.67 | 34.51 ±5.28 | 33.38 ±6.75 | 22.94 ±6.39 | 34.79 ±6.21 | 39.17 ±13.84 | 32.21 ±9.49 | 0.103 |
| *a*<sup>*</sup> | 18.26 ±6.64 | 7.43 ±0.33 | 14.01 ±5.99 | 10.79 ±5.96 | 10.33 ±4.70 | 13.12 ±4.61 | 6.97 ±7.66 | 11.25 ±4.93 | 0.190 |
| *b*<sup>*</sup> | 11.69 ±7.58<sup>ab</sup> | -2.15 ±7.58<sup>b</sup> | 14.54 ±3.98<sup>ab</sup> | 13.40 ±8.87<sup>ab</sup> | 2.18 ±7.06<sup>ab</sup> | 15.89 ±5.11<sup>ab</sup> | 16.11 ±10.08<sup>a</sup> | 10.3 ±6.82<sup>ab</sup> | 0.023 |
| Δ*E*<sup>*</sup> | - | 20.99 | 5.74 | 7.80 | 15.30 | 7.23 | 14.13 | 7.15 |  |
| Abdomen |  |  |  |  |  |  |  |  |  |
| *L*<sup>*</sup> | 41.99 ±7.35 | 35.42 ±2.21 | 44.85 ±9.18 | 43.06 ±7.05 | 33.61 ±6.26 | 37.69 ±6.70 | 45.54 ±12.11 | 32.95 ±7.69 | 0.294 |
| *a*<sup>*</sup> | 32.18 ±8.52 | 20.88 ±5.43 | 24.64 ±11.47 | 21.00 ±12.44 | 23.57 ±1.11 | 25.47 ±3.30 | 16.46 ±9.78 | 21.01 ±6.37 | 0.453 |
| *b*<sup>*</sup> | 22.21 ±8.94 | 13.40 ±9.01 | 24.85 ±8.91 | 24.76 ±11.01 | 14.22 ±5.105 | 20.85 ±0.42 | 24.56 ±9.53 | 15.42 ±8.10 | 0.434 |
| Δ*E*<sup>*</sup> | - | 15.76 | 8.49 | 11.51 | 14.43 | 8.08 | 16.29 | 15.89 |  |

*L*<sup>\*</sup>, Lightness; -*a*<sup>\*</sup>, greenness; +*a*<sup>\*</sup>, redness; - *b*<sup>\*</sup>, blueness; + *b*<sup>\*</sup>, yellowness. $\Delta E^*$ values represent colour differences between the treatment and the control group (C). C: Control; T: *Tetraselmis suecica*; D: *Dunaliella salina*; N: *Nannochloropsis* sp. \*Kruskal-Wallis test. Different superscripts on the same row represent there is a statistical difference (P<0.05).

### 3.3. Interactions of pseudo-green water mixture on overall performance

The larval morphometric table and mixture designs provide a visualisation of the contribution effect of the individual algal species towards the animal’s length, weight, carapace length and colouration (Figure 1, Figure 2). The largest total lengths resulted from a mixture of *D. salina* and *Nannochloropsis* sp., skewed towards *Nannochloropsis* sp. The largest mean weights and carapace lengths were obtained from mixtures comprised of *T. suecica, D. salina* and *Nannochloropsis* sp., and *T. suecica and Nannochloropsis* sp., respectively. The monocultures show less impact, with the shortest mean carapace lengths and lowest mean weights occurring with the *T. suecica* monoculture (TS) and control treatments. With lowering levels of *Nannochloropsis* sp. within mixtures, lengths and weights appear to reduce, and combinations of *T. suecica* and *D. salina* result in relatively lower levels of growth. From the ANOVA test, *Nannochloropsis* sp. was the only contributor of significant influence (p < 0.05). Still, results imply it is most effective as part of a mixture (Figure 1), which is reflected in the research for other species. For example, dried *Nannochloropsis* sp. mixed with *Isochrysis* sp. produced an improved feed conversion ratio (FCR) and a greater muscle proportion of fatty acids eicosapentaenoic acid and arachidonic acid when fed to the studied Atlantic cod (*Gadus morhua*) at 15 % dietary replacement (Walker & Berlinsky, 2011). More pertinently for lobster larvae, several benefits are reported from the resultant studied species when used as an application for zooplankton enrichment. In one study, researchers enriched rotifers with *Chlorella vulgaris* and *Nannochloropsis oculata* at a 50:50 ratio and reported an increase in stress resistance and improved growth in the barramundi (*Lates calcarifer*) larvae (Thepot et al., 2016). Since lobster larvae do not actively hunt their prey in early stages, instead relying on encounters stimulated by chemoattraction and, to a certain degree, chance encounters (Kurmaly et al., 1990), adding mixed microalgae could benefit growth and performance by providing a constant supply of more easily accessible nutrients.

**Figure 1:**
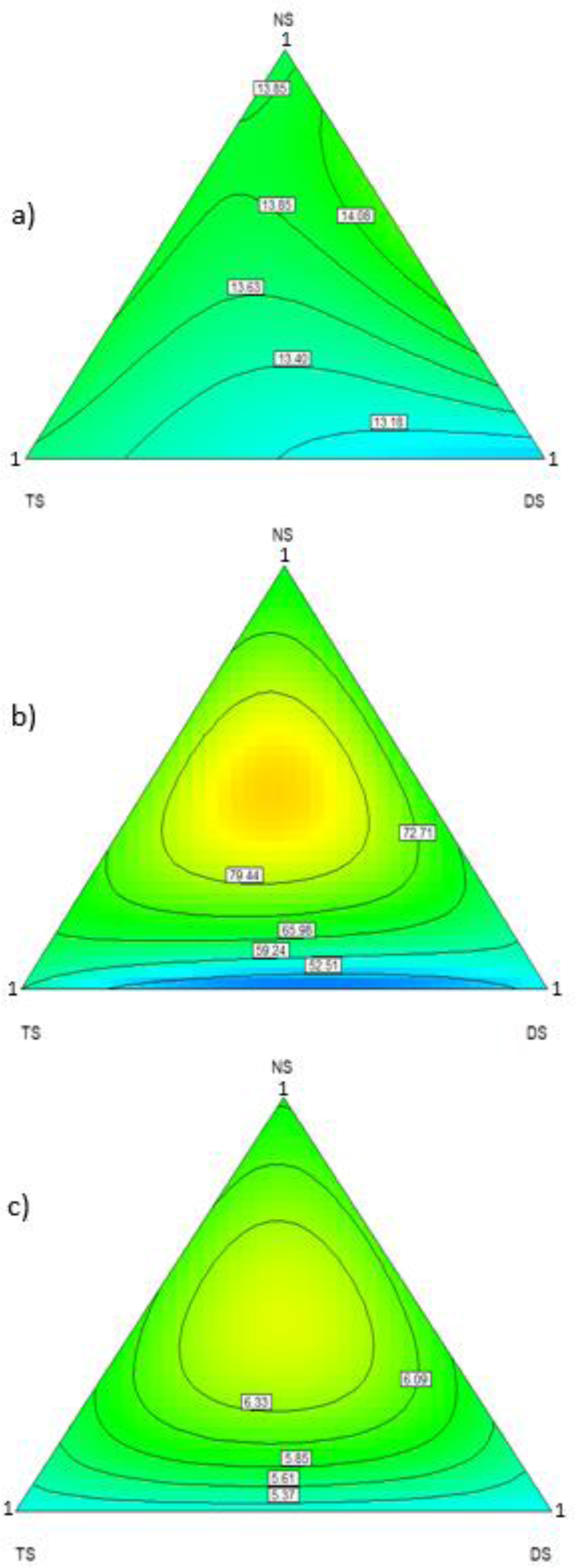
Various mixture designs showing influence of individual algal species on day 13 in relation to a) Total length (mm), b) Mean weight (mg), c) Mean carapace length (mm) measurements (TS: *Tetraselmis suecica;* DS: *Dunaliella salina;* NS: *Nannochloropsis* sp.).

**Figure 2:**
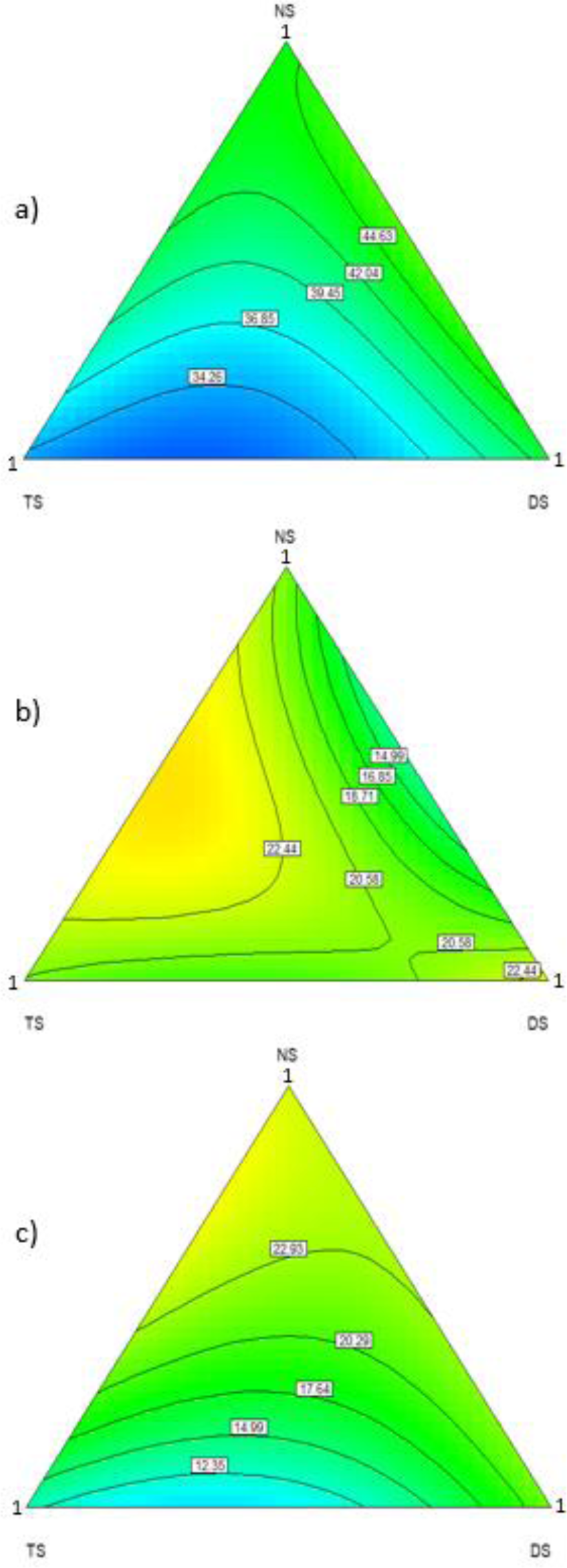
Various mixture designs showing the influence of individual algal species on day 13 in relation to mean colour metrics of the claw, cephalothorax, and abdomen. a) L*, Lightness; b) a*, +redness/-greenness; c) b*+yellowness/-blueness. TS: *Tetraselmis suecica;* DS: *Dunaliella salina;* NS: *Nannochloropsis* sp.).

The L*a*b* colour mixture designs (Figure 3) display the mean colour metrics of the claw, cephalothorax, and abdomen in relation to a) L*, Lightness; b) a*, +redness/-greenness; c) b*+yellowness/-blueness and the various microalgae species studied. Lightness appears to be skewed more towards mixtures of *Dunaliella salina and Nannochloropsis* sp. Redness appears to be more prominent in mixtures containing *T. suecica and Nannochloropsis* sp., and yellowness appears to be more applicable to mixtures containing all three studied species of microalgae *T. suecica, D. salina* and *Nannochloropsis* sp., with a slight skew towards *Nannochloropsis* sp. These results also suggest that colour metrics are better managed in mixtures containing all three studied microalgaes. This appears to be logical as each microalgae contains varying levels of carotenoid pigments e.g., *Dunaliella salina* is known for containing a significant amount of β-carotene, *Nannochloropsis* sp., contains a large amount of violaxanthin and T. suecica has been shown to contain a variety of carotenoids including α-carotene, β-carotene and their derivatives (Anusree et al., 2023; Di Lena et al., 2019; Ren et al., 2021). ANOVA for the cubic mixture models (Table 4) displayed no significant variation between the mixtures carried out in this study in relation to morphometric and L*a*b colouration parameters.

**Table 4:** The interaction of different algal species on European lobster (*Homarus gammarus*) larval mean morphometric development and shell colouration.

| Parameter | Source | F value | P value |
| --- | --- | --- | --- |
| Total length | Model | 0.69 | 0.657 |
|  | NC/TS | 0.14 | 0.709 |
|  | NC/DS | 2.09 | 0.163 |
|  | TS/DS | 0.02 | 0.886 |
|  | NC/TS/DS | 0.26 | 0.613 |
| Weight | Model | 1.84 | 0.154 |
|  | NC/TS | 1.10 | 0.310 |
|  | NC/DS | 0.78 | 0.389 |
|  | TS/DS | 0.83 | 0.376 |
|  | NC/TS/DS | 2.72 | 0.118 |
| Carapace length | Model | 2.34 | 0.070 |
|  | NC/TS | 1.85 | 0.188 |
|  | NC/DS | 1.42 | 0.247 |
|  | TS/DS | 0.02 | 0.867 |
|  | NC/TS/DS | 1.98 | 0.175 |
| $L^*$ | Model | 1.44 | 0.264 |
|  | NC/TS | 0.18 | 0.675 |
|  | NC/DS | 0.46 | 0.506 |
|  | TS/DS | 0.89 | 0.360 |
|  | NC/TS/DS | 0.23 | 0.637 |
| $a^*$ | Model | 0.77 | 0.603 |
|  | NC/TS | 0.55 | 0.471 |
|  | NC/DS | 3.03 | 0.102 |
|  | TS/DS | 0.21 | 0.656 |
|  | NC/TS/DS | 0.58 | 0.459 |
| $b^*$ | Model | 1.54 | 0.232 |
|  | NC/TS | 0.78 | 0.390 |
|  | NC/DS | 1.81 | 0.966 |
|  | TS/DS | 1.37 | 0.259 |
|  | NC/TS/DS | 4.28 | 0.998 |

$L^*$ , Lightness; $a^*$ , +redness/-greenness; $b^*$ +yellowness/-blueness. TS: *Tetraselmis suecica*; DS:
*Dunaliella salina*; NS: *Nannochloropsis* sp.

The present study noted the highest survival ratios in the mixed *Dunaliella salina and Nannochloropsis* sp., (D + N) and the mixed *Dunaliella salina, Nannochloropsis* sp., and Tetraselmis sp., (T + D + N) treatments, each achieving a 20.00 % survival rate. The lowest recorded survival rate of 12.5 % was obtained in the monoalgal *Nannochloropsis* sp., (N) treatment, which equated to a 60 % reduction in survival rate compared to the D + N and T + D + N treatments. The techniques used in this study were, therefore, insufficient to surpass the planktonic larval phase bottleneck of 20% (Addison and Bannister, 1994; Daniels et al., 2010; Ellis et al., 2015; Goncalves et al., 2023; Powell et al., 2017).

It is difficult to isolate origins of mortality given the number of causative effects: cannibalism, natural life-history strategy, inadequate nutrition, disease, genetic issues, and feed being unsuitable for stage IV larvae (Goncalves, 2021; Goncalves et al., 2022; Kurmaly et al., 1990; Nguyen et al., 2018; Pechenik, 1999; Romano & Zeng, 2017). Research shows that feed quality determines survival outcomes in controlled environments (Schoo et al., 2014). Survival rates obtained in this study indicate that utilising pseudo-green water in rearing had no detrimental effects on the survival of European lobster larvae. Therefore, there is room for improvement in this area of research, and a focused study on enhancing survival through pseudo and green water techniques is needed. Feeding on conspecifics has been proven to greatly improve survival outcomes in European lobster larvae (Powell et al., 2017), but this would be unsuitable for scaling up hatchery production.

Interpreting the causes of these results is difficult without comprehensive proximate analyses on both algae and larvae. Though the causative effects of the algal diets can be speculated upon based on existing nutrient approximations from established studies, interpretations must be considered preliminary. For example, even when algae are cultured under standard conditions, nutrition profiles are highly variable (Brown et al., 1997; Enright et al., 1986; Guedes & Malcata, 2012). Therefore, determining diet macronutrient content is essential before assigning any estimate of feed quality (Whyte et al., 1990). Proximate analyses on post-trial lobster specimens would also be needed to compare and correlate nutrition profiles, similar to current techniques employed in other species, e.g., whiteleg shrimp (*Penaeus vannamei*, Barreto et al., 2023).

Using proximate analyses of juvenile or mature European lobster would not be suitable for interpreting the relationship between larval growth and feed nutrition content found in this study (Yeap et al., 2022). Goncalves (2021) has demonstrated that European lobster nutrition requirements vary significantly throughout their life history, with post-metamorphoses juveniles increasing their dietary carbohydrate requirements compared to the more lipid-focused larval stages.

## 4. Conclusion

Pseudo-green water techniques remain a novel area of research in producing European lobster larvae. Pseudo-green water techniques contrast with green water techniques and provide algal enrichment during the most critical life stage development for European lobster, the larval stages only. Unlike clear water techniques, the pseudo-green water technique provides the larvae with lipids, β-carotene, and other important nutrients for enrichment and development. It aids in achieving an elevated survival rate within hatcheries. However, the limited knowledge of the nutritional requirements of European lobster larvae remains a considerable obstacle to hatchery production for restocking, stock enhancement, and future farming operations. Nevertheless, indirect feed nutritional enhancement through adding microalgae has proven to be an effective larval-rearing technique, growth and increased survival outcomes in multiple crustaceans and fish species. This study identified that combining a mixture of microalgae to the rearing environment can increase growth performance indicators and the survival rates of European lobster larvae. However, it is noted that there is a disparity between algal species; therefore, further investigations to screen other mixture designs are required. Pseudo-green water techniques may be the next step in lobster rearing within hatchery programs for stock enhancement and aquaculture.

## CRediT

**MEB:** Formal analysis; Validation; Visualisation; Roles/Writing - original draft; Writing - review & editing.

**JG:** Conceptualisation, Data curation; Formal analysis; Investigation; Methodology; Validation; Visualisation; Roles/Writing - original draft; Writing - review & editing.

**ERF**: Investigation; Methodology; Validation; Writing - review & editing.

**JH**: Conceptualisation; Visualisation; Roles/Writing - original draft; Writing - review & editing.

**SJD:** Visualisation; Roles/Writing - original draft; Writing - review & editing.

**SM**: Roles/Writing - original draft; Writing - review & editing.

**AHLW**: Conceptualisation; Data curation; Formal analysis; Funding acquisition; Investigation; Methodology; Project administration; Resources; Software; Supervision; Validation; Visualisation; Roles/Writing - original draft; Writing - review & editing.

